# Patterns of association between mothers and offspring and their outcomes in a polygynous ungulate

**DOI:** 10.64898/2026.05.07.723517

**Authors:** Adam Z. Hasik, Natasha Robinson, Fiona E. Guinness, Sean Morris, Alison Morris, Tim Clutton-Brock, Josephine M. Pemberton

## Abstract

Prolonged association between mothers and their offspring is common in ungulates, with the level of maternal investment likely to play a central role in shaping this trait. Here we examined patterns of association between mothers and offspring over time, the apparent benefits of association to offspring, and costs to mothers. We analyzed 40 years’ worth of census data from an individually-monitored, food-limited population of red deer (*Cervus elaphus*) on the Isle of Rum, Scotland. Starting from birth, female calves associated more frequently with their mothers than male calves in their first year. Calves also associated less with their mothers if the mother did not conceive a new calf. Association frequency decreased with mother’s age and population density, and survival over the first year was not related to mother-calf association. Yearlings, now in their second year, were more often associated with their mothers if they were female, if there was no subsequent calf (or the subsequent calf died as a neonate), and if they were still being suckled. Increased association between mothers and yearlings was associated with increased survival to adulthood at 28 months, but suckling a yearling did not improve its probability of survival. For individuals that reached maturity, increased association in the yearling year was associated with slightly shorter adult life spans. The level of association between a calf and mother was not associated with the mother’s immediate survival or fecundity. Our findings suggest that juveniles born to poor-condition mothers benefit from prolonged association through improved yearling survival.

## Introduction

Traits spread through populations when they are associated with increased fitness, and parental care traits are selected for when they benefit the parent’s overall fitness (Clutton-Brock, 1991; Williams, 1966). One way that parents can increase their lifetime fitness is by enhancing offspring survival and reproduction (Alonso-Alverez and Velando, 2012; Hamilton, 1964). While parental care traits enhance offspring fitness, they are often also associated with costs, such as reduced adult survival and reproductive success (Alonso-Alverez and Velando, 2012; Clutton-Brock, 1991; Ewer, 1968). Given the nature of these trade-offs, the benefits to the offspring must outweigh the costs to the parents for traits to be favored (Balshine, 2012; Clutton-Brock, 1991; Klug et al., 2012). Parents are selected to vary their offspring investment based on changes in the corresponding payoffs; if parental costs outweigh offspring benefits, then the care trait will reduce (Alonso-Alverez and Velando, 2012; Higashi and Yamamura, 1993; Trivers, 1972; Williams, 1966).

Parental investment in one offspring may not only affect a parent’s own survival but also reduce the investment in other offspring (Clutton-Brock, 1991; Trivers, 1972; Williams, 1966). This is particularly evident in long-lived iteroparous species, where parents face investment trade-offs between current and future offspring over their entire reproductive lifespan (Clutton-Brock et al., 1996; Hamel et al., 2011; Martin and Festa-Blanchet, 2010). Parental care traits generally benefit new offspring more than previous offspring, leading to reduced or ceased care for older offspring (Clutton-Brock, 1991). When no new reproduction occurs, mothers can maintain care for their existing offspring and enhance their fitness (Clutton-Brock et al., 1982b).

Parental investment also varies with offspring life history demands, for example sex differences shaped by sexual selection (Clutton-Brock, 1991; Trivers, 1972). In polygynous systems, where males depend on resources, particularly early in life, for growth and mating success, mothers play a crucial role in enhancing their sons’ reproductive success (Clutton-Brock et al., 1979; Festa-Bianchet et al., 2000; Reiter et al., 1978). In contrast, the reproductive success of daughters is less strongly-associated with early maternal input, leading to stronger maternal investment in males early on, interpreted to be a strategy to maximize offspring fitness (Clutton-Brock et al., 1982b; Kruuk et al., 1999; Maynard Smith, 1980). Such differences are reflected in the relative costs to mothers of rearing male versus female offspring (Clutton-Brock et al., 1981; Froy et al., 2016; Gomendio et al., 1990).

In species with polygyny and female philopatry, as in many ungulates, the lack of female dispersal allows mothers to enhance the fitness of female offspring later in life. For ungulates in particular, protracted association (which is the product of both the mother and calf) is often regarded as a form of parental care as daughters may benefit from protection and access to resources including reduced displacement from preferred forage (Andres et al., 2013; Brookshier and Fairbanks, 2003; Clutton-Brock, 1991; Green et al., 1989; Holand et al., 2012; Testa, 2004). While continued association may increase fitness for an offspring, it may also come at a cost to a mother if it increases competitions for resources (Charest Castro et al., 2018; Clutton-Brock et al., 1982a; Higashi and Yamamura, 1993; Stewart et al., 2005).

Parental investment is also expected to vary with the energy state of the parent. An extensive literature now documents that in natural populations performance in many traits increases in young animals and later senesces (Jones et al., 2014), including in aspects of maternal investment such as offspring birth date and weight (Hayward et al., 2015; Nussey et al., 2009). Broadly, parental care might be expected to decline with parental age, though it could also manifest as extended care without further reproduction (Clutton-Brock, 1984).

Similarly, ecological conditions can drive variation in care by adjusting the payoffs involved (Clutton-Brock et al., 1996; Hamel et al., 2010; Lawrence, 1990; Rowell, 1991). At high population density, increased intraspecific competition may limit resource access and affect fitness traits and components such as growth and reproduction (Siepielski et al., 2020; Stewart et al., 2005). Mothers may therefore allocate resources by prioritizing their own survival and reproduction and reducing investment in offspring (Festa-Bianchet et al., 2019; Martin and Festa-Blanchet, 2010) and care might be expected to be density-dependent.

Here, we investigate the dynamics of mother-offspring association in the polygynous, female-philopatric red deer (*Cervus elaphus*) over the first two years of an offspring’s life. We used 40 years’ worth of census data from the Isle of Rum to analyze factors associated with the extent of association, as well as the potential fitness benefits and costs of association. Association is not a precise measure of care in that it involves no obvious resource transfer and it is the result of both the mother and the offspring’s behavior. Nevertheless, we assume that mothers and offspring would not associate unless they both have an interest in the outcome.

In the study population, females are philopatric and tend to occupy the home ranges of their mothers while males leave their mother’s home range within two to four years of birth (Clutton-Brock et al., 1982b). Females typically have their first calf in their third or fourth year of life and then continue to produce a single calf most years until death. Rearing a calf is costly, as demonstrated by the fact that females are less likely to survive or conceive after rearing a calf, especially a male (Clutton-Brock et al., 1983; Froy et al., 2016; Guiness et al., 1979). In red deer, conception is strongly condition-dependent (Albery et al., 2021; Albon et al., 1983; Albon et al., 1986; Hasik et al., 2026), and in the Rum population females do not produce a calf (i.e., are true yeld) about one year in three on average. If a female fails to conceive, this sets up an opportunity for enhanced association with her previous calf, including the possibility of suckling it into a second year. Prolonged lactation with delayed weaning enhances offspring fitness in other mammals (i.e., northern elephant seals, Reiter et al., 1978), and previous analyses of the study population showed that association provides fitness benefits to red deer calves through increased resource access and reduced aggression (Clutton-Brock et al., 1982b). One motivation for this research is that current open shooting seasons for red deer in the UK begin when a calf is 4-5 months old. Best practice is to avoid orphaning a calf since this lowers its probability of survival (Andres et al., 2013). Our study documents the natural progression of mother-calf association and likely precision with which mother-calf pairs can be identified and shot.

In this study, our objectives were to provide an overview of the pattern of mother-offspring association over the first two years of life, and then to investigate three questions. First, how does mother-offspring association vary in relation to calf sex, the mother’s reproductive status and age, and population density? Second, how does association relate to juvenile survival? Third, how does association in the calf’s early life relates to a mother’s immediate survival and fecundity?

## Materials and Methods

### Study population and data collection

We collected data for this study from a population of individually-recognized red deer in the north block of the Isle of Rum, Scotland (57°N,6°20’W), a study described in detail in Clutton-Brock et al. (1982b) and Pemberton et al. (2022). In brief, Rum has a wet, mild climate and contains a mixture of high-quality (i.e., nutritious) grassland and low-quality (i.e., high in tannins, less nutritious) blanket bog and heath. The study area runs ∼4km north to south and ∼3km east to west, totaling ∼12.7km^2^. The deer are unmanaged within the study area though risk being shot when they wander outside it, as there are no fences. The “deer year” begins with the calving season at the start of May and ends at the end of April of the next year. Around 90% of calves born in the study area are caught soon after birth and permanently marked; natural markings are also used for individual recognition. Since 1974 regular censusing of the deer has occurred five times per month covering most months of the year (mainly excluding in the calving and rutting seasons), providing data on most deer living in the study area. Records are made of individual deer, their locations, and their group membership. Detailed life history data is collected during the calving and rutting seasons, as well as during mortality searches over winter. Most calves are weaned by January. However, some continue to take milk for over a year. This ‘yearling suckling’ has been systematically recorded since 1985.

### Data overview and measurements

For this study we used *n* = 98,311 observations of *n* = 2,742 individual juveniles over 40 years. Observations of these juveniles were made over 1,540 censuses: the first being in May 1985 and the most recent in July 2025. Each census record contained the individual’s ID, sex, age, location (to the nearest 100 m) and group ID. Individuals were considered to be in the same group when they were within 50m of their nearest group member, a value derived from the investigation of neighbor distances (Clutton-Brock et al., 1982b). When juveniles were recorded multiple times in one census, we only included the first observation to avoid artificially-inflating the association frequency. To avoid scenarios where the juvenile or mother may have been obscured rather than genuinely missing from a group, we only included a record when the juvenile and its mother were seen in the same census, either in the same group or in different groups. We excluded juveniles without a known birth date (mostly because the mother was outside the study area when she gave birth), those that died at less than 139 days (mostly neonatally), or those that were shot from the study.

We categorized a juvenile as either associating with its mother when they were seen in the same group (1) or non-associated when seen without her (0, as in Guinness et al., 1979). We conducted most statistical analyses with the raw data (i.e., a binary variable). However, when required as a predictor variable, we calculated an association index by dividing the number of observations when a juvenile was seen associated by the total number of observations of that juvenile within the period investigated (i.e., a proportion).

We categorized juvenile data into two age groups, with age measured in days between the juvenile’s birth date and the date of census. “Calves” were in their first year of life and the data consisted of census observations when individuals were under 365 days old, while “yearlings” were in their second year and the data were collected when they were between 365 and 730 days old.

To understand the factors affecting mother-calf association we considered calf sex (either female, *n* = 1,378, or male, *n* = 1,364), age (calf or yearling), mother’s age and age squared (to test for non-linear relationships), mother’s reproductive status, and population density. For yearlings we included yearling suckling status, a binary factor denoting if a juvenile suckled as a yearling (i.e., during its second year of life). The mother’s reproductive status value differed for calves and yearlings. For calves, the mother could be pregnant (having conceived in the rut when the calf was approximately four months old) or not pregnant. For a yearling, a mother could be in one of four categories during the yearling year: true yeld: did not become pregnant, so no follow-on calf, summer yeld: had a calf but it died before October (often as a neonate), winter yeld: had a follow-on calf but it died in the winter, milk: had a calf and raised it through its first year of life. Population density was an annual measure of the number of females aged one year or older regularly using the study area.

Second, we investigated the relationship between mother-offspring association (either during the calf or yearling year) and offspring survival. We defined two survival metrics for both the calves and yearlings: survival to the end of the year (1) or not (0) and adult lifespan defined as the number of days between 28 months and death. We excluded all deer without recorded death dates (i.e., they were still alive or death was not recorded, principally because they had emigrated from the study area) from lifespan analyses. We also excluded individuals that were shot as they ranged outside the study area from survival analyses when needed (i.e., an individual shot at age three years or more could still be in the analysis of survival as a yearling).

Third, we investigated the relationship between mother-offspring association in the first year of a calf’s life and the mother’s overwinter survival (to April 30^th^) and subsequent fecundity.

### Statistical analysis

#### How does mother-offspring association vary in relation to calf sex, mother’s reproductive status and age, and population density?

We conducted all analyses in R v.4.2.3 (R Core Team 2025). For all of the models below we standardized all continuous variables to have a mean of 0 and SD of 1. Our first test of mother-juvenile association used all records from calves in their first year of life (*n* = 50,589 records from *n* = 2,285 individuals from *n* = 1,484 censuses). We constructed a mixed-effects logistic regression with calf sex, maternal age, maternal age^2^, pregnancy status, and population density as fixed effects. Given that during January-May a mother may be investing resources in a growing fetus, we also fit a two-level time-of-year factor with “1” denoting the first seven months of a calf’s life, June to December, and “2” denoting the second five months, January to May, as well as the interaction between this timing and pregnancy status. Calf ID, mother ID, and year were included as random effects to control for repeated measures of the calf, maternal effects, and annual variation, respectively.

Our second test of juvenile association used records for yearlings (*n* = 34,489 records from *n* = 1,688 individuals from *n* = 1,531 censuses). We constructed a similar model to that of the calf association test, with yearling sex, maternal age, maternal age^2^, mother’s reproductive status, population density, and suckling status as fixed effects with yearling ID, mother ID, and year as random effects. As noted above, here reproductive status was a four-level factor according to the mother’s putative reproductive effort in the current year.

### How does association relate to juvenile survival?

To understand the potential benefits of association we investigated relationships between association and annual survival (short-term benefit) and lifetime (long-term benefit). For calves (*n* = 716 individuals) we used survival to the end of the first year as our response variable in a mixed-effects logistic regression with the association index for each calf in their first year, calf sex, maternal age, maternal age^2^, pregnancy status, and population density as fixed effects with mother ID and year as random effects. For yearlings (*n* = 300 individuals) we wanted to understand if association during the second year of life affected survival into adulthood, thus we constructed a model similar to that of the calves but included suckling or not as a fixed effect. Note that our yearling analysis only included the deer that survived their first year of life and made it to the yearling age class.

To understand if association during the early years had long-term benefits to lifespan, we constructed a Gaussian mixed-effects model with lifespan as the response variable (log-transformed to approximate normality), the association index for each calf in their first year and calf sex as fixed effects, with mother ID and year as random effects. We repeated this analysis with the yearling data, using a similar model structure but including whether the yearling was suckling or not as a binary fixed effect.

### How does association in the calf ’s early life relates to a mother’s immediate survival and fecundity?

To understand the relationships between mother-calf association and the survival and fecundity of the mother we focused on the first six months of a calf’s life, providing insight into whether the first few months of association came at a cost to the mother before the expression of survival or gestation costs. We built two models, one where we used the mother’s overwinter survival as the response variable with calf sex and association index as fixed effects in a mixed-effect logistic regression model with year and individual ID as random effects. In the second model we used fecundity (binary: 1 for females that had a calf in the subsequent year and 0 for females that did not) as the response variable in a mixed-effect logistic regression with calf sex, the association index, population density, age, and age^2^ as fixed effects with year and individual ID as random effects. We included additional factors in the fecundity model as all of these variables are known to affect fitness (both survival and fecundity) in this system, but we could not include them in the survival model as survival was very high in these data and including the additional variables resulted in convergence issues for the model.

## Results

### How does mother-offspring association vary in relation to calf sex, mother’s reproductive status and age, and population density?

Association between a mother and her offspring varied with the time of year, offspring sex, and offspring age (Fig. 1a). Typically, starting from birth mother-calf association for both sexes was high throughout the calf’s first year, with female calves on average more closely-associated than males. Association dropped markedly in both sexes at age 1, especially males, and there was an additional precipitous drop in males, from which association never fully recovered, when they reached 15-16 months at the time of the rut. For males this coincided with aggression directed at yearlings by harem-holding stags.

**Fig. 1.**
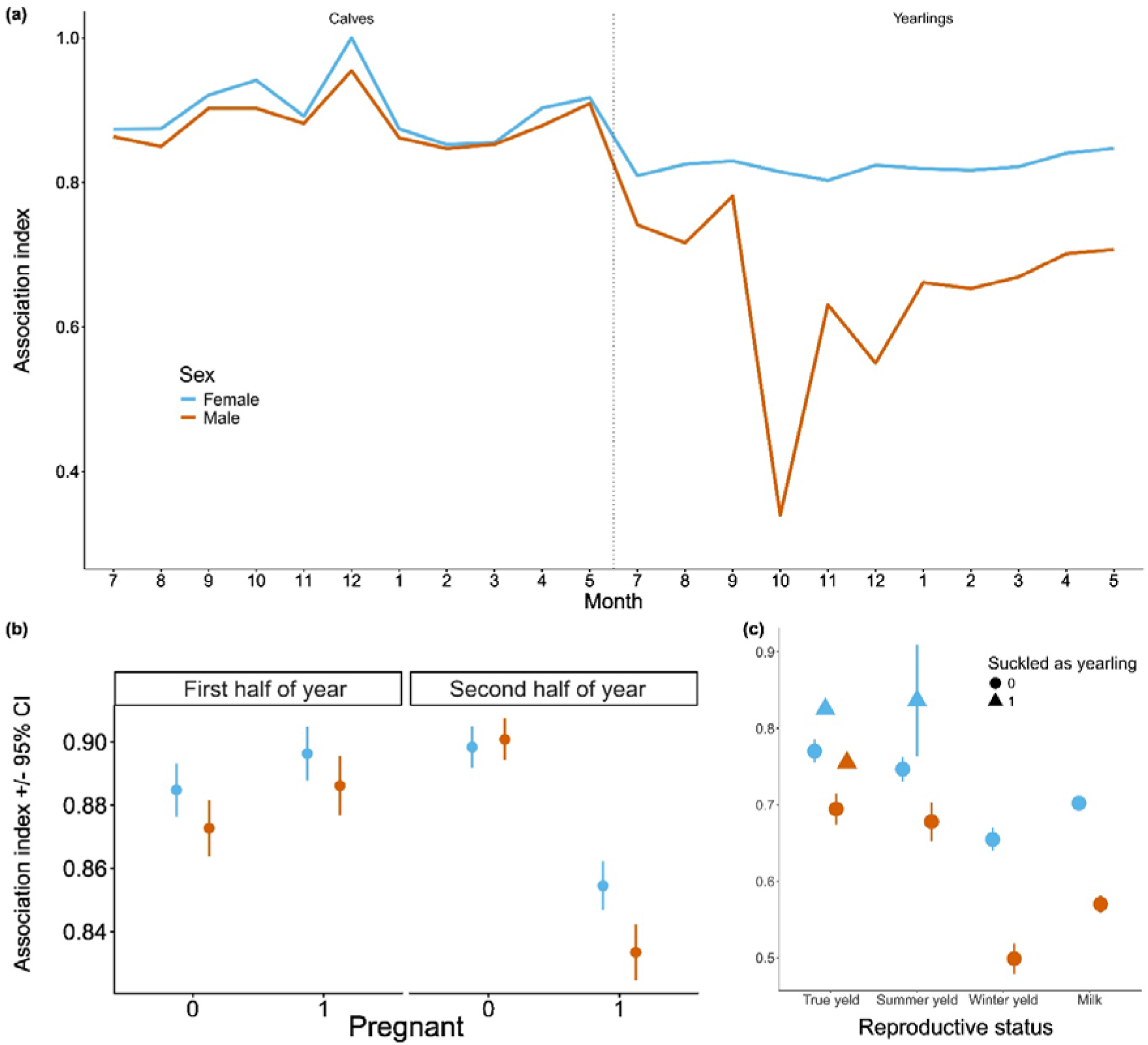
Mother-offspring association varied with offspring age, offspring sex, and the reproductive status of the mother. (a) shows the mean association trends in calves and yearlings from all observations across the first two years of life, with the dotted line denoting the start of the new deer year and the point at which a calf becomes a yearling. Note that “6” is absent from the x-axis as we do not census in June due to the calving season. (b) shows the association index for calves on the y-axis with the mother’s pregnancy status on the x-axis, with separate panels for each half of the first (calf) year. (c) shows the association index for yearlings on the y-axis with the mother’s reproductive status on the x-axis. Color in all panels denotes offspring sex, while error bars in (b) and (c) denote the 95% CI’s.

Our model relating association to calf sex, mother’s pregnancy status, annual timing, and population density revealed a significant interaction between annual timing and pregnancy status (Χ*^2^_1_* = 163.45, *p* < 0.001). From June to December association was lower with male calves than female calves (Χ*^2^_1_* = 6.33, *p* = 0.01, Fig. 1b) but maternal pregnancy status did not impact association (Χ*^2^_1_* = 3.71, *p* = 0.05, Fig. 1b). Association was also lower for older mothers (correlation coefficient = −0.30, Χ*^2^_1_* = 4.24, *p* = 0.04), but there were no relationships with maternal age^2^ (Χ*^2^_1_* = 3.73, *p* = 0.05) or annual population density (Χ*^2^_1_* = 1.81, *p* = 0.18).

From January to May the sex of the calf did not relate to the association index (Χ*^2^_1_* = 0.76, *p* = 0.38, Fig. 1b). Association decreased with maternal age (correlation coefficient = −0.36, Χ*^2^* = 4.42, *p* = 0.04), but was not related to maternal age^2^ ( *^2^* = 1.51, *p* = 0.22). Association was lower if the mother was pregnant than if she was not (Χ*^2^_1_* = 33.32, *p* < 0.001) and decreased with annual population density (correlation coefficient = −0.17, Χ*^2^_1_* = 9.34, *p* = 0.002).

In our model of yearling association, association varied with a yearling’s sex and mother’s reproductive status, as expected in a male-biased dispersal system. We found lower association in male yearlings than females (Χ*^2^* = 195.80, *p* < 0.0001, Fig. 1c) and that the association varied with the level of investment into a subsequent calf (effect of mother’s reproductive status, Χ*^2^_3_* = 56.85, *p* < 0.0001, Fig. 1c), with lower assocation when the mother was rearing a new calf. As might be expected, yearlings that suckled associated more with their mothers than those did not suckle (Χ*^2^_1_* = 62.14, *p* < 0.0001, Fig. 1c). There was no relationship with maternal age (Χ*^2^* = 0.38, *p* = 0.54), or maternal age^2^ ( *^2^* = 0.01, *p* = 0.95), or with population density (Χ*^2^_1_* = 1.43, *p* = 0.23),

### How does association relate to juvenile survival?

In calves we found that, as previously-demonstrated, males had reduced survival over the first year of life compared to females (Χ*^2^_1_* = 19.02, *p* < 0.0001). Perhaps surprisingly, increased association with the mother was not associated with survival (Χ*^2^_1_* = 0.02, *p* = 0.90; Fig. 2a), but calves whose mothers were pregnant again had strongly reduced survival (correlation coefficient = −0.70, Χ*^2^_1_* = 10.18, *p* = 0.002). In this particular model, first-year survival was not related to birth year density (Χ*^2^_1_* = 0.25, *p* = 0.62), maternal age (Χ*^2^_1_* = 1.46, *p* = 0.41), or maternal age^2^ (Χ*^2^_1_* = 1.57, *p* = 0.21).

**Fig. 2.**
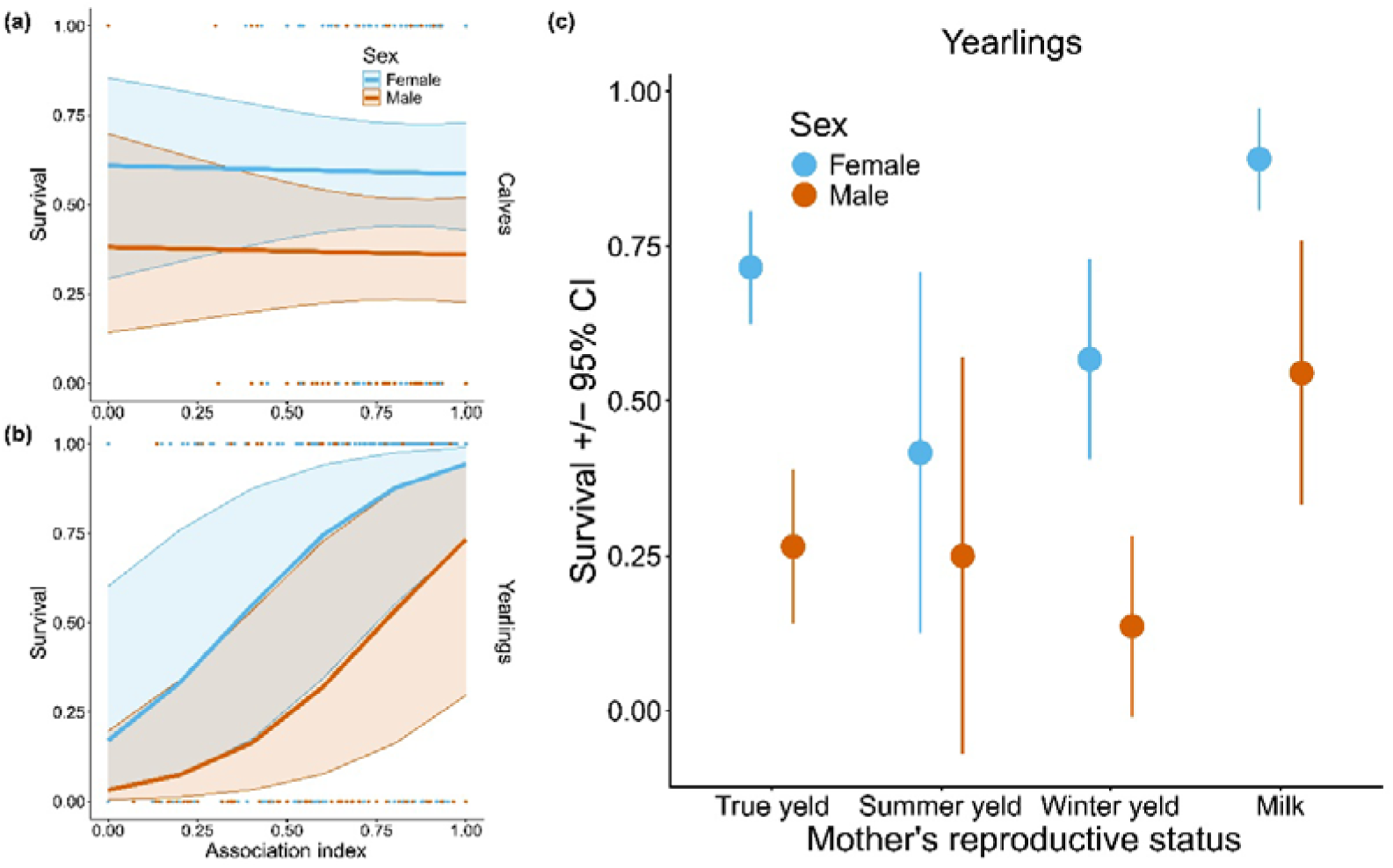
Survival to age 1 in calves (a) and age 2 in yearlings (b), with survival on the y-axes, the association index on the x-axes. Regressions in both panels are model-derived likelihoods of survival across the range of observed association indices, with bands representing 95% CI’s. Points in both plots are observed data from individual deer, and color denotes offspring sex. (c) shows the association index for yearlings on the y-axis with the mother’s reproductive status on the x-axis. Color in all panels denotes offspring sex, while error bars in (c) denote the 95% CI’s.

In yearlings, males were again less likely to survive to adulthood (Χ*^2^_1_* = 25.96, *p* < 0.0001). We also found that yearling survival to adulthood increased with the association index (Χ*^2^_1_* = 24.95, *p* < 0.0001, Fig. 2b) and varied as a function of mother’s reproductive status (Χ*^2^_3_* = 15.21, *p* = 0.002, Fig. 2c), with yearlings whose mothers raised their new calves through their first year of life (milk hinds) having the highest survival. Yearling survival to adulthood related to mother’s age both linearly (correlation coefficient = 2.07, Χ*^2^* = 4.70, *p* = 0.03) and quadratically (correlation coefficient = −2.33, Χ*^2^_1_* = 4.37, *p* = 0.04) but there was no association with density. Conditional on association, suckling as a yearling was not related to survival (Χ*^2^_1_* = 2.11, *p* = 0.15), nor did it explain variation in survival when fit without association.

With regard to lifespan after 28 months, our analysis revealed that the positive benefits of increasing association did not extend beyond the juvenile stage. We found that increased association during the first year of life was not associated with longer lifespans in our analysis of the calves (Χ*^2^_1_* = 1.83, *p* = 0.18, Fig. 3a), but we found that increased association during the second year of life was associated with shorter lifespans (Χ*^2^_1_* = 4.72, *p* = 0.03, Fig. 3b). Specifically, we found overall lifespan differences of 1.4 and 1.38 years for females and males, respectively, across the gradient of association index values. We also found that males had shorter lifespans than females, in both our calf (Χ*^2^_1_* = 29.25, *p* < 0.0001) and yearling (Χ*^2^_1_* = 24.04, *p* < 0.0001) models, though there was no relationship between suckling status and lifespan for yearlings (Χ*^2^_1_* = 0.01, *p* = 0.92).

**Fig. 3.**
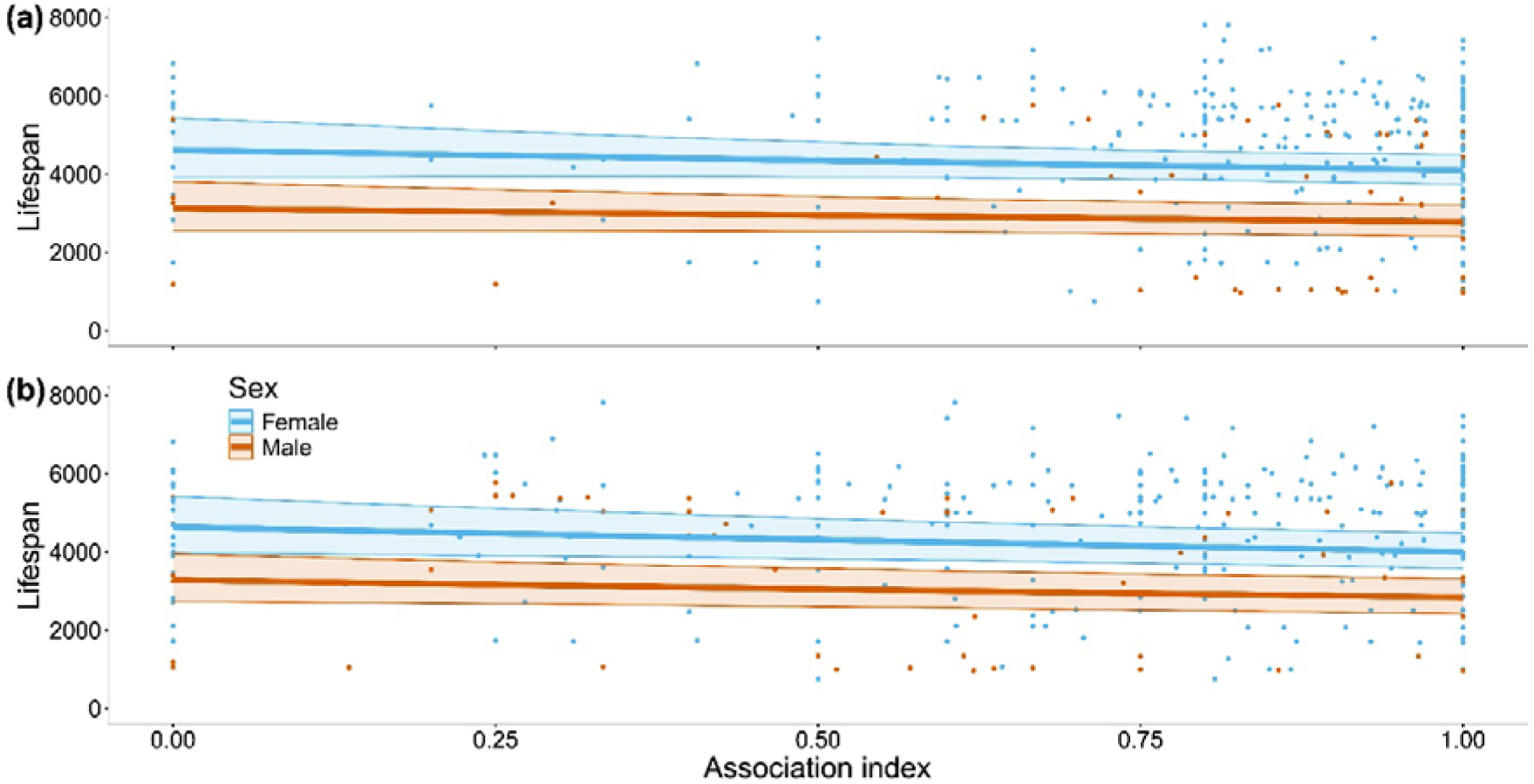
Lifespan was not related to mother-calf association (a) but it was for mother-yearling association (b). The y-axes show lifespan in days, the x-axes show the association index. Regressions in both panels are model-derived estimates of lifespan across the range of observed association indices, with bands representing 95% CI’s. Points in both plots are observed data from individual deer, and color denotes offspring sex.

### How does association in the calf ’s early life relates to a mother’s immediate survival and fecundity?

We found no evidence that increased association with a calf affected the immediate fitness of the mother. Increased association with the calf over the first 6 months was not associated with decreased survival (Χ*^2^* = 1.21, *p* = 0.27, Fig. 4a) or decreased fecundity (Χ*^2^* = 2.13, *p* = 0.14, Fig. 4b). Female survival was lower for mothers that raised a male calf (Χ*^2^_1_* = 7.02, *p* = 0.01, Fig. 4a), but there was no relation with fecundity (Χ*^2^_1_* = 0.04, *p* = 0.85, Fig. 4b). Fecundity declined with population density (correlation coefficient = −0.31, Χ*^2^_1_* = 5.21, *p* = 0.02), and we also found evidence for linear (correlation coefficient = 2.37, Χ*^2^_1_* = 12.06, *p* = 0.001) and quadratic (correlation coefficient = −3.04, Χ*^2^_1_* = 15.63, *p* < 0.0001) relationships between age and fecundity.

**Fig. 4.**
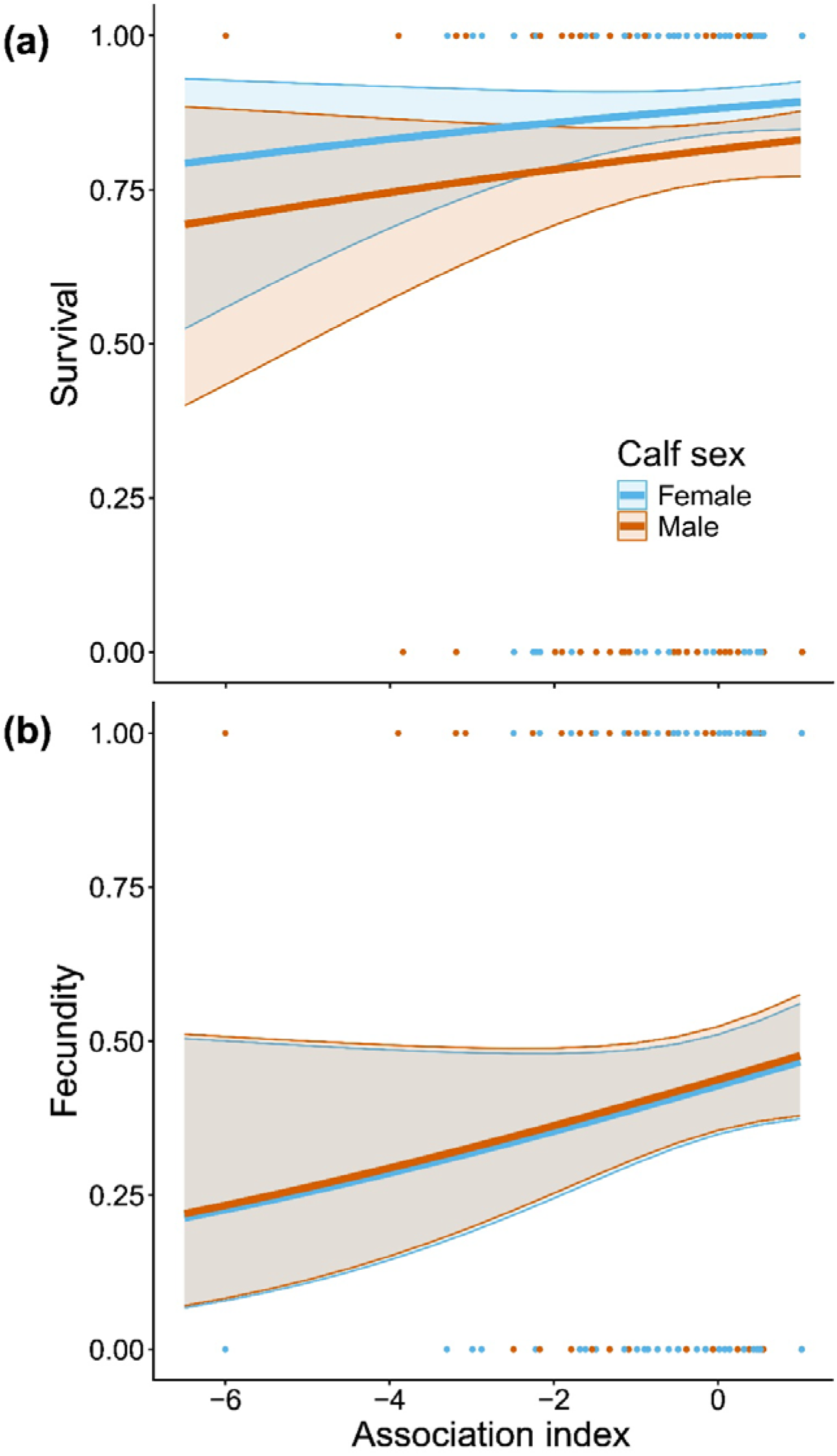
Female survival (a) and fecundity (b) did not vary as a function of association with calves. The y-axes show survival (a) and fecundity (b), while the x-axes show the association index over the first six months of the calf’s life. Regressions in both panels are model-derived estimates of the probability of survival (a) and fecundity (b) across the range of observed association indices, with bands representing 95% CI’s. Points in both plots are observed data from individual deer, and color denotes offspring sex.

## Discussion

Using 40 years’ worth of data from an exceptionally well-characterized red deer study system we investigated the predictors and apparent consequences of mother-offspring association over the first two years of a juvenile’s life. On average, mothers and calves of both sexes were seen together on between 80% and 100% of censuses over the first year (Figure 1a).

When surviving offspring reached one year old, mother-offspring association dropped to about 80% for females and much lower for males for whom the rut, when they were about 16 months old, was associated with a sharp decline in association, which recovered to about 70% for the rest of the year (Figure 1a). We now discuss our detailed analyses of mother-calf association, in which a recurring theme is that the condition of a mother, as revealed by whether she conceived a follow-on calf, impacts many downstream events.

### How does mother-offspring association vary in relation to calf sex, mother’s reproductive status and age, and population density?

Mother-offspring association across the first two years of life varied with the reproductive status of mothers. Association declined in the second part of the calf year if a female became pregnant with a follow-on calf (Figure 1b). This decreased association, also found in Guinness et al. (1979), is associated with weaning the existing calf in the face of nurturing the new embryo and preparing for post-parturition costs (Clutton-Brock et al., 1983).

Presumably, later stages of pregnancy become too costly for mothers to continue investing in previous young, leading to a reduction in association (Clutton-Brock, 1991; Klug and Bonsall, 2014).

Association between mothers and yearlings was strongly associated with whether there was a follow-on calf and its fate (Figure 1c). Mother-yearling association was lowest if the mother had a follow-on calf that either did or did not survive the winter (milk or winter yeld, respectively) and was higher if the mother did not produce a follow-on calf (true yeld) or the follow-on calf died neonatally (summer yeld). As with the second part of the calf year, investing in previous offspring while caring for a newborn would increase a mother’s burden and risk jeopardizing their newborns’ success (Clutton-Brock et al., 1982b). Evidence of prioritization of newborns has been shown in other ungulate species, with some instances of mothers actively driving yearlings away (Charest Castro et al., 2018; Hall, 2023; Hirth, 1977; Rowell, 1991), a behavior also seen in the Rum deer. For females without a follow-on calf, maintaining association with a yearling, potentially improving its survival and future reproduction, becomes the optimal strategy for enhancing their overall fitness (Brookshier and Fairbanks, 2003; Charest Castro et al., 2018; Green et al., 1989; Testa, 2004). Yearling suckling was almost exclusively observed in true yeld mothers, and as might be expected, such yearlings were more often seen with their mothers (Figure 1c).

Consistent with previous work (Guinness et al., 1979), we found only modest evidence for sex differences in mother-offspring association within the first year of the calves’ lives, when both sexes are largely dependent on their mothers (Clutton-Brock et al., 1982a). In the first seven months of life, males were slightly less often associated with their mothers than females, but this was not true for the remaining five months (Figure 1a, b). This may be related to greater exploratory behavior by male calves or disruption associated with the rut. Sex differences in association developed in the second year of life, when female yearlings showed much higher association with their mothers (Figure 1c). The difference emerged as the rut approached (Figure 1a), and male yearling association declined sharply around the rut, as adult stags aggressively drive yearling males out of harems (Clutton-Brock et al., 1979; Guinness et al., 1979) (Figure 1a). Male yearling association never fully recovered after this period. This aligns with the well-characterized social system of red deer, in which philopatric females remain in natal groups while males gradually disperse (Clutton-Brock, 1991).

Guinness et al. (1979) proposed that increased male dispersal may result from inherently exploratory nature of males or maternal aggression. Male yearlings may also be less-inclined to maintain a close association with their mothers if they yield fewer benefits from prolonging the relationship. In orphaned calves, maternal loss is associated with reduced daughter survival over multiple years, while it was only associated with first year survival in sons, implying males gain minimal benefits from maternal presence later in life (Andres et al., 2013). In contrast to the limited advantages for males to remaining associated, numerous studies have demonstrated that females in social, philopatric species continue to benefit throughout their lives by staying close to their mothers (Brookshier and Fairbanks, 2003;

Green et al., 1989; L’Heureux et al., 1995; van Noordwijk et al., 2012). An alternative hypothesis is that maternal aggression triggers male dispersal, which could explain some of the sex-dependency in association rates (Guinness et al., 1979). Males can increase their lifetime reproductive success by excelling in growth, a trait closely linked to their maternally-supported early development (Clutton-Brock et al., 1982b). In contrast, females do not gain as many fitness benefits from enhanced early growth and do not pay as large a cost for poor early development (Festa-Bianchet et al., 2000). Given males are more costly to rear than females, mothers may prompt their dispersal as a means to avoid further investment (Berube et al., 1996).

Association between mothers and calves in their first year was influenced by mother’s age. Association declined with mother’s age in both the first seven months and the second five months of the calf’s life. Two possible and non-mutually exclusive processes may lie behind this observation: as females age, they may be in gradually-worsening condition (e.g., Albery et al., 2022; Albery et al., 2024), leading them to care less for their calves.

Alternatively, as mothers get older and more experienced, they are less attentive and more relaxed about letting their calf wander off or somehow instill more confidence in their calf to wander off. Previous studies of ageing effects in wild animal populations, including the Rum deer, generally show an increase in performance in early life and a decline in late life, identified by fitting a quadratic term in models (Jones et al., 2008; Nussey et al., 2009). Age^2^ was not associated with mother-calf association in our models, suggesting that the second explanation may be more likely. Mother-yearling association was not related to the mother’s age or age^2^, indicating that any maternal age effects dissipate after an offspring’s first year.

Mother-calf association was unrelated to population density in the first half of the calf year, but was negatively-associated with density in the second part of the calf year. This suggests that when conditions are tough, mothers and calves are more likely to separate, perhaps to forage separately. This is consistent with both the prediction that an increase in competition would reduce levels of association (Clutton-Brock et al., 1987b; Johnson, 1986; Martin and Festa-Blanchet, 2010) and the well-documented density-mediated fitness costs in this population (Albon et al., 1987; Clutton-Brock et al., 1987a; Clutton-Brock et al., 1987b).

Namely, increasing density reduces fitness directly through increased competition for limited resources and indirectly through increased exposure to parasites (Hasik et al., 2025a; Hasik et al., 2026), with increasing parasite burden further reducing fitness in juvenile and adult deer (Acerini et al., 2022; Hasik et al., 2025b). In contrast, mother-yearling association showed no relationship with density.

### How does association relate to juvenile survival?

Calf survival over the first winter was not related to mother-calf association, although as previously-documented, males had lower survival that females (Figure 2a). This result is perhaps a little surprising given weaning and the reduction in association if a mother becomes pregnant again. A possible explanation lies in variation in maternal condition. If a mother is in sufficient condition to conceive a follow-on calf, then presumably she was in good condition when provisioning the focal calf, such that it has a high probability of survival despite early weaning and reduced association. Conversely, a calf whose mother does not conceive a follow-on calf may be in poorer condition and may not wean the focal calf as quickly as a pregnant female, promoting higher association. It seems possible these processes could lead to similar outcomes for pregnant and non-pregnant mothers in terms of calf survival.

In contrast to calf survival, yearling survival was strongly related to mother-yearling association, again with male survival lower than female survival (Figure 2b). In the same statistical model, survival of both sexes was highest for offspring of milk mothers, intermediate for offspring of true yeld mothers and lowest in winter yeld mothers (Figure 2c), suggesting another role for the mother’s condition: a mother that rears two calves in succession is clearly in better condition than a mother that didn’t have a follow-on calf or one that lost a follow-on calf in winter. After accounting for association, a yearling that was still being suckled did not have improved survival compared to one that was not. Yearling suckling is tightly bound with the failure of the mother to conceive a follow-on calf (true yeld; Figure 1c). Our interpretation of this relationship is that suckling a yearling enables a poor-condition mother to compensate a poor-condition yearling to the extent that its survival matches that of yearlings that are not suckled.

In an interesting parallel to the calf and yearling survival results, adult lifespan was not associated with mother-calf association but it *was* with mother-yearling association, though in an unexpected direction: higher mother-yearling association was associated with *shorter* adult lifespan (Figure 3a,b). Our hypothesis here is that we are observing a syndrome in which mother-yearling association is highest in mothers in poorest condition in the offspring’s first year (no follow-on calf, consequently true yeld; Figure 1c) and this is associated with improved yearling survival, but the effects of a poor first year are nevertheless carried over into adult lifespan. This would be consistent with previously-documented early life impacts on adult performance in the study population (Albon et al., 1987; Kruuk et al., 1999).

### How does association in the calf ’s early life relates to a mother’s immediate survival and fecundity?

Mother-calf association in the first six months of the calf’s life did not predict the mother’s survival or fecundity over the following year. Although the relative costs of gestation and lactation have been extensively documented in the Rum deer (Albery et al., 2021; Clutton-Brock et al., 1989; Froy et al., 2016), our results are not surprising, because by definition all mothers in our analysis had gestated and reared a calf over its first 139 days (i.e., into the autumn). This restriction ensured that calves that died neonatally or over summer (meaning that they were rarely seen in censuses) did not enter the data set, but it does mean all mothers in the analysis had successfully reared a calf into the autumn and are not differentiated according to likely costs up to that point in time.

One motivation for this study is to inform deer managers about the extent to which mothers and offspring are associated and its consequences. We have previously shown that the death of the mother in the first year of an offspring’s life reduces the probability that a calf survives (Andres et al., 2013). Therefore, managers try to shoot calves at the same time as their mothers, but the success with which they can do this depends of their ability to identify the mother-calf pair. Our analyses suggest that mother-calf pairs are reliably seen together in the first part of the female shooting season in the UK, from October to December, but less reliably together in the second part of the season in January-February, suggesting that shooting early in the season would give greater accuracy. A further advantage of shooting early in the season is that the animals are in better condition and have not yet undergone the privations of winter.

Our study was conducted using decades-worth of high-quality data from an exceptionally well-studied population of red deer, yet it is not without its limitations. We have focused on association between mothers and offspring, which is not the same as the actual transfer of resources involved in parental investment, and is also a consequence of the behavior of two different individuals. Nevertheless, we make the assumption that two individuals would not associate unless there was some advantage to one or both of them. Milk transfer is one of the obvious modes of maternal care. Unfortunately, in red deer suckling occurs only a few times a day: even in newborn calve there are only 2-5 bouts per day (Clutton-Brock et al., 1982b). Consequently, it is hard to tell precisely when weaning occurs for a given mother-offspring pair. Yearling suckling is only detected in spring and especially during the calving season (May-June) when there are many observers in the study area. Other benefits that probably accrue from association are knowledge of forage and shelter locations in the landscape and protection of the offspring from harassment by other individuals.

In conclusion, we have shown that in a food-limited red deer population, mother-offspring association over the first two years is strongly determined by whether or not the mother has a follow-on calf. Since conception of a follow-on calf is condition-dependent (Albon et al., 1983; Albon et al., 1986), our results are consistent with a finely-tuned maternal response in which failure to conceive a new calf results in continued association with the previous calf which in turns improves the survival prospects of that offspring.

## Author contributions

AZH and JP conceived the study. AZH, NR, and JP designed the study and analyses. FG, SM, AM, TCB, and JP collected field data. AZH and NR performed modelling work and analyzed data. AZH wrote the first draft of the manuscript, and all authors contributed substantially to revisions.

## Acknowledgements

We thank NatureScot for permission to work on the Isle of Rum and the many volunteers and researchers who have helped at the field site during this study. This work was supported by the Leverhulme Trust (RPG 2022-220) to JMP and AZH. AZH benefitted from the musical inspiration of Bloc Party.

